# Protein secondary structure determines the temporal relationship between folding and disulfide formation

**DOI:** 10.1101/564740

**Authors:** Philip J Robinson, Shingo Kanemura, Xiaofei Cao, Neil J Bulleid

## Abstract

How and when disulfides form in proteins during their folding is a fundamental question in cell biology. Two models describe the relationship between disulfide formation and folding, the folded precursor model, in which formation of nascent structure occurs prior to the disulfides and the quasi-stochastic model where disulfides form prior to complete domain folding. Here we investigate oxidative folding within a cellular milieu of three structurally diverse substrates in order to understand the folding mechanisms required to achieve correct cysteine coupling. We use a eukaryotic translation system in which we can manipulate the redox conditions and produce stalled translation intermediates representative of different stages of translocation. We identify different disulfide bonded isomers by non-reducing SDS-PAGE. Using this approach, we determined whether each substrate followed a folding driven or disulfide driven mechanism. Our results demonstrate that the folding model is substrate-dependent with disulfides forming prior to complete domain folding in a domain lacking secondary structure, whereas disulfide formation was absent at this stage in proteins with defined structural elements. In addition, we demonstrate the presence and rearrangement of non-native disulfides specifically in substrates following the quasi-stochastic model. These findings demonstrate why non-native disulfides are prevented from forming in proteins with well-defined secondary structure.

**Significance statement:** A third of human proteins contain structural elements called disulfide bonds that are often crucial for stability and function. Disulfides form between cysteines in the specialised environment of the endoplasmic reticulum (ER), during the complex process of protein folding. Many proteins contain multiple cysteines that can potentially form correct or incorrect cysteine pairings. To investigate how correct disulfide pairs are formed in a biological context, we developed an experimental approach to assess disulfide formation and rearrangement as proteins enter the ER. We found that a protein domain with atypical secondary structure undergoes disulfide orchestrated folding as it enters the ER and is prone to incorrect disulfide formation. In contrast, proteins with defined secondary structure form folding dependent, native disulfides. These findings show how different mechanisms of disulfide formation can be rationalised from structural features of the folding domains.

## Introduction

A central question in protein folding is how proteins that contain multiple cysteines achieve the correct disulfide pattern. Intramolecular disulfide bonds consist of covalent cross-links between cysteines within a polypeptide (1), and their formation imposes conformational restraints that influence protein folding, stability and function. Disulfides form through an oxidative reaction between paired cysteines in the presence of an appropriate catalyst (2). In eukaryotic cells, the endoplasmic reticulum (ER) is the main site for disulfide formation, it provides a favourable redox environment and catalysts in the form of resident protein disulfide isomerase (PDI) family members (3). When polypeptides contain more than two cysteines, correct pairing is required to form the native disulfide pattern; the spatial positioning of cysteine sidechains is required for this correct pairing, closely linking disulfide formation to the process of conformational folding.

Two models have been proposed that broadly explain the relationship between disulfide formation and conformational folding:(i) the folded precursor mechanism, in which nascent structure forms first, which then positions paired cysteines favourably to form a disulfide and (ii) the quasi-stochastic model, in which cysteines in an unfolded precursor pair more randomly, and the resulting disulfide influences further folding (4). Recent studies have identified substrates that favour the folded precursor mechanism (5–7), while other studies have identified contributions from both mechanisms at different stages of the folding process (8). Incorrect (non-native) disulfides can also form during folding, which are subsequently reduced and re-arranged (9, 10). In the ER, this reduction step is performed by specific PDI family members (11, 12) using reducing equivalents that originate in the cytosol (13). Evidence shows that non-native disulfides can act as important intermediates in the native folding of certain proteins (10).

Folding and disulfide formation in the cell is a vectorial process. During the early stages of translation, nascent polypeptides enter the ER through the Sec-translocon (14). The space constraints of the ribosome/Sec-complex limit structure formation until enough polypeptide has entered the ER-lumen (15). The addition of each amino acid to the polypeptide expands the repertoire of intra-molecular interactions and the diversity of potential folds that might form. Both native and non-native disulfides can form at this co-translational stage (16, 17). The exposure of the nascent polypeptide to the ER, as well as the folding of the nascent polypeptide and the accessibility of PDI family members dictates native disulfide formation (18). Non-native disulfides may also form owing to the partial exposure of nascent polypeptides to the ER, which prevents completion of folding, leading to incorrect cysteine coupling.

The question of how proteins with multiple cysteines achieve the correct disulfide pattern is a difficult one to address due to the structural diversity of disulfide-containing proteins and the technical barriers that hinder folding assays performed in a biologically relevant context. In an earlier study, we devised an approach to investigate the folding of a protein containing a single disulfide as it emerges into the ER lumen (7). Here, we expand on this approach to study nascent polypeptides with diverse structures and complex disulfide patterns to investigate the relationship between protein folding and correct disulfide formation. Our results demonstrate the influence of protein secondary structure on the fidelity of disulfide formation within a cellular context.

## Results

### Experimental design

The three proteins used in this study, β2-microglobulin (β2M), prolactin and the disintegrin domain of ADAM10, were chosen for their diversity of secondary structure and disulfide bonding (Fig. 1A). We added a C-terminal extension to each protein (Fig. 1B) to act as a tether to the ribosome and to enable the folding domain to enter the ER lumen before completion of translation (Fig 1 Ci). DNA templates were synthesised that encode specific polypeptide lengths that lack stop codons. The templates were transcribed and the resulting mRNA used to programme translation reactions that produce stalled nascent chains of differing lengths. Translations were performed using a rabbit reticulocyte lysate supplemented with semi-permeabilised (SP) cells to study folding in the ER (19) or without SP cells to represent folding in the cytosol. All samples were treated with NEM on completion to irreversibly modify thiols and freeze the disulfide status of the samples for downstream processing.

**Figure 1.**
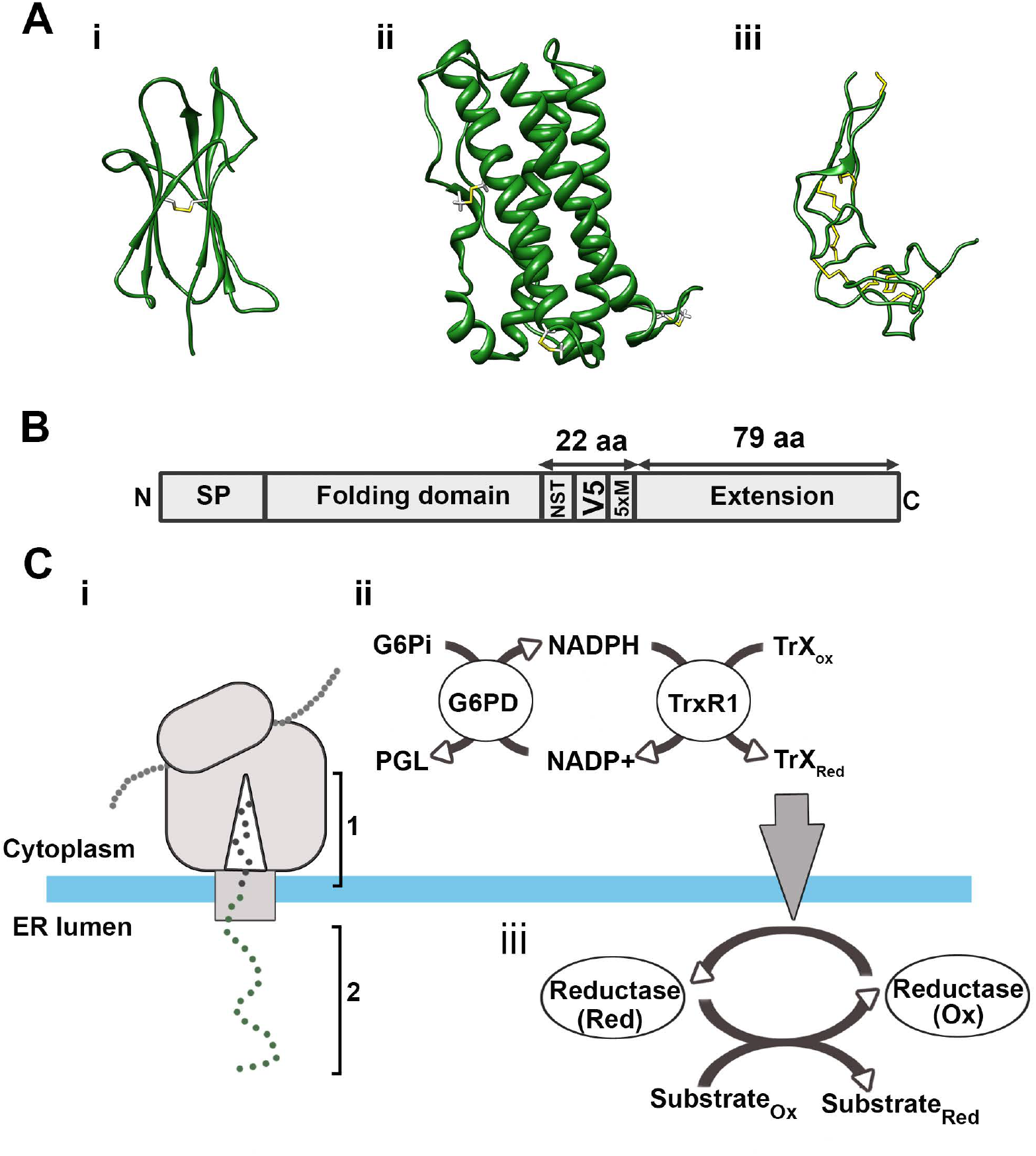
An experimental system to monitor disulfide formation and rearrangement in translation intermediates. (A) Ribbon diagrams representing the substrates used in this study: (i) human β2M (PDB 1A1M); (ii) human prolactin (PDB 1RW5); and (iii) the disintegrin domain of human ADAM10 (PDB 5L0Q). Disulfide bonds are highlighted in yellow. (B) Schematic illustrating the C-terminal extension added to each folding domain to produce the extended constructs. The position of the signal peptide (SP), glycosylation site (NST), V5 epitope and five methionine residues (5xM) are indicated. (C) (i) Schematic of the ribosome-Sec complex expressing an ER-exposed nascent polypeptide. Phase 1 shows how the C-terminal extension retains the polypeptide attached to the ribosome to allow Phase 2. In Phase 2 the N-terminal folding domain of the polypeptide is fully exposed to the ER lumen. (ii) G6Pi drives the cytosolic reducing pathway, which (iii) is the source of reducing equivalents for disulfide rearrangements in the ER.

We adjusted the redox status of the translation reactions by adding specific components to the rabbit reticulocyte lysate. In all, we used three different lysates in our experiments. (1) A reducing lysate supplemented with DTT, in which proteins synthesised are unable to form disulfides in the absence or presence of SP-cells. (2) An oxidising lysate that contained no additional components; this lysate allows disulfides to form in proteins synthesised in the absence or presence of SP cells. And (3) a redox-balanced lysate supplemented with G6Pi to drive G6PD and TrxR1 activity (13) (Fig.1 Cii); this lysate is sufficiently reducing to prevent disulfide formation in proteins synthesised without SP cells but allows disulfide formation in translocated proteins when SP-cells are present (13). Proteins synthesised in the redox-balanced lysate supplemented with SP cells have the capacity for disulfide reduction and rearrangements. This is because the activity of cytosolic G6PD and TrxR1 is the source of reducing equivalents for an ER-reductive pathway, which is required to reduce non-native disulfides (Fig. 1 Ciii). Thus, by comparing disulfide formation in oxidising and redox-balanced lysates, we can assess the influence of the reducing pathway on the fidelity of disulfide formation.

### Stochastic disulfide formation is absent in β2M

β2M is a small, β-sheet rich protein with two cysteines and a single disulfide (Fig. 1Ai). It has an N-terminal signal peptide that is cleaved following ER-targeting (Fig. 2 Ai). We can follow disulfide formation in β2M by non-reducing SDS-PAGE as it has a faster mobility than the reduced protein (7). When wild type β2M was translated without SP cells in the redox-balanced or reducing lysates, the resulting product migrated as a single species corresponding to the reduced preprotein (Fig., 2Aii, lanes 1 and 2; pre_red_). When translated in the oxidising lysate a faster migrating species was detected indicating disulfide formation (lane 3; pre_ox_). When β2M was synthesised in the presence of SP cells, ER-targeting resulted in signal peptide cleavage with the translation product from the reducing lysate migrating as two species representing reduced preprotein (pre_red_) and reduced mature protein (mat_red_) (lane 4). The translation product from the redox-balanced and oxidised lysates also migrated as two species but the mature protein (mat_ox_) runs faster relative to mat_red_ indicating disulfide formation. Disulfide formation also occurred in preprotein translated in the oxidising lysate (lane 6) as seen in the absence of SP-cells. These results demonstrate that disulfide formation takes place following polypeptide translocation into the ER in both redox-balanced and oxidising lysates.

**Figure 2.**
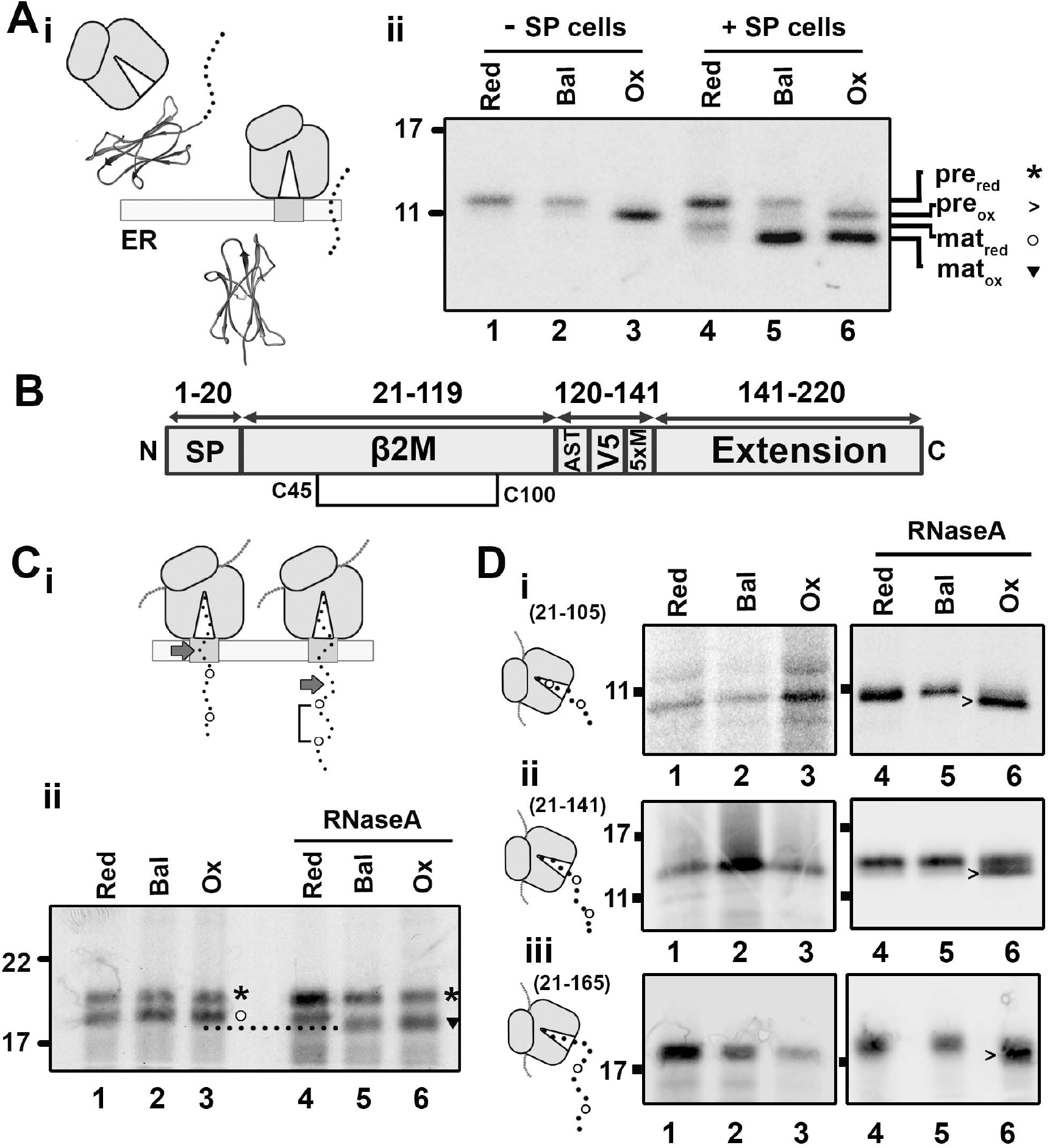
β2M-disulfide formation is folding driven. (A) (i) Diagram illustrating signal peptide cleavage that occurs following translocation of β2M (ii) Non-reducing SDS-PAGE of immunoisolated wild type β2M translated with or without SP cells in reducing (Red), redox-balanced lysate (Bal) or oxidising (Ox) lysate. The migration of preprotein (pre) and mature protein (mat) when oxidised (ox) or reduced (red) is indicated with associated symbols (*,>,o, triangle) (B) Organisation of the extended β2M construct with the single disulfide (C45-C100) displayed. (C) (i) Diagram illustrating the dependence of disulfide formation on domain exposure, arrows highlight the end of the β2M folding domain. (ii) Non-reducing SDS-PAGE shows isolated 175 intermediates translated with SP cells in redox-balanced (Bal), oxidising (Ox), and reducing (Red) lysate. RNaseA treatment following translation is indicated. (D) Non-reducing SDS-PAGE analysis of isolated ribosome/nascent chain complexes representing β2M (i) 21-105, (ii) 21-141, and (iii) 21-165 translated in redox-balanced (Bal), oxidising (Ox) or reducing (Red) lysate without SP cells. Samples treated with RnaseA following translation are indicated. Schematics next to each panel indicate the predicted cysteine exposure at each intermediate length. All gels in this figure are representative of at least 3 repeats.

In a previous study we used an extended-β2M sequence (Fig. 2B) to investigate folding events during ER entry and found that disulfide formation is a folding-driven process that requires the β2M sequence to be fully translocated into the ER (Fig. 2Ci) (7). This finding is consistent with a structured precursor mechanism of folding. We performed these experiments in a redox-balanced lysate and consequently the ER reductive pathway may prevent any disulfide formation prior to translocation of the folding domain. To investigate if this was the case we translated the 175 intermediate of extended-β2M in either the absence (oxidising lysate) or presence (redox-balanced lysate) of a robust ER reductive pathway (Fig. 2 Cii). At this intermediate length, both cysteine residues will have exited the Sec-complex while part of the folding domain remains within. The RNA transcript did not contain a stop codon thereby preventing its release from the ribosome. The resulting translation products had the same migration independent of the lysate used (lanes 1-3) indicating a lack of any disulfide formation when the nascent chain is tethered to the ribosome. Only upon release of the polypeptide from the ribosome following RNase treatment did disulfides form, specifically in samples translated in oxidising and redox-balanced lysates (triangle) (lanes 5 and 6). This result demonstrates that the lack of disulfide formation prior to full translocation of the β2M folding domain is independent of the ER reductive pathway. This indicates that β2M follows the structured-precursor mechanism for disulfide formation despite redox conditions that are more likely to favour disulfide formation.

To investigate the role of ER factors in folding and disulfide formation we synthesised stalled RNCs of β2M in the oxidising lysate in the absence of SP-cells. We previously showed that these conditions are sufficient to induce disulfide formation in wild type β2M released from the ribosome (Fig 2Aii). Polypeptide synthesis in the absence of ER factors to regulate disulfide formation may favour stochastic disulfide formation. For these experiments, we used a β2M sequence that did not contain an amino-terminal signal sequence in case this affected protein folding. Three intermediate lengths were chosen based on estimates assuming that ~40 amino acids span the length of the ribosome tunnel (2/3 of the amino acids required to span the entire ribosome/Sec-complex) (7, 20). The schematics in Fig. 2D illustrate the predicted cysteine exposure in each case. Following translation in the different lysates, stalled RNCs were either isolated by ultracentrifugation through a sucrose cushion or immunoisolated following RNaseA treatment (Fig, 2D i, ii, iii). No disulfide formation occurred for stalled and released intermediates translated in redox-balanced or reducing lysates (lanes 1, 2, 4, 5). The stalled intermediates translated in the oxidising lysate showed no evidence of disulfide formation either (lanes 3), despite exposure of both cysteines to the oxidising lysate in the 141 and 165 intermediates. Disulfide formation occurred only following release of the nascent chains from the ribosome (lanes 6).

These results with β2M support our earlier findings (7) that suggested disulfide formation requires a folding-driven mechanism. Here we extend this conclusion to include conditions where stochastic disulfide formation is more likely. The translations in oxidising lysate both with and without SP cells show that the entire folding domain needs to exit the ribosome before disulfide formation can take place. This suggests that stochastic disulfide formation does not occur during β2M folding and that disulfides form only once the polypeptide folds.

### Disulfide formation in prolactin favours a structured precursor-folding model

The next substrate studied was bovine prolactin, which has an α-helical structure and three native disulfide bonds (21). A schematic illustrates the extended-prolactin construct (Fig. 3A). We monitored N-linked glycosylation at position 230 to determine the intermediate length required for prolactin domain exposure to the ER lumen (Fig 3B). Following translation of increasing lengths of the extended-prolactin, glycosylation occurred from an intermediate length of 305 amino acids (Fig. 3Bi). We calculated that 75 amino acids (305–230) are required to span the distance between the P-site of the ribosome and the active site of the oligosaccharyl transferase (OST). Taking into account the ~12 residues it takes to reach from the exit of the Sec-complex to the OST, we estimate that 63 residues span the ribosome/Sec-complex (Fig. 3B ii), the same value observed with the β2M construct (7). Based on this result, we predicted the length of polypeptide exposed to the ER for each intermediate as shown in both table and topology-diagram (Fig 3. Biii). For the remaining experiments with prolactin, we used a construct without the glycosylation site to simplify SDS-PAGE analysis.

**Figure 3.**
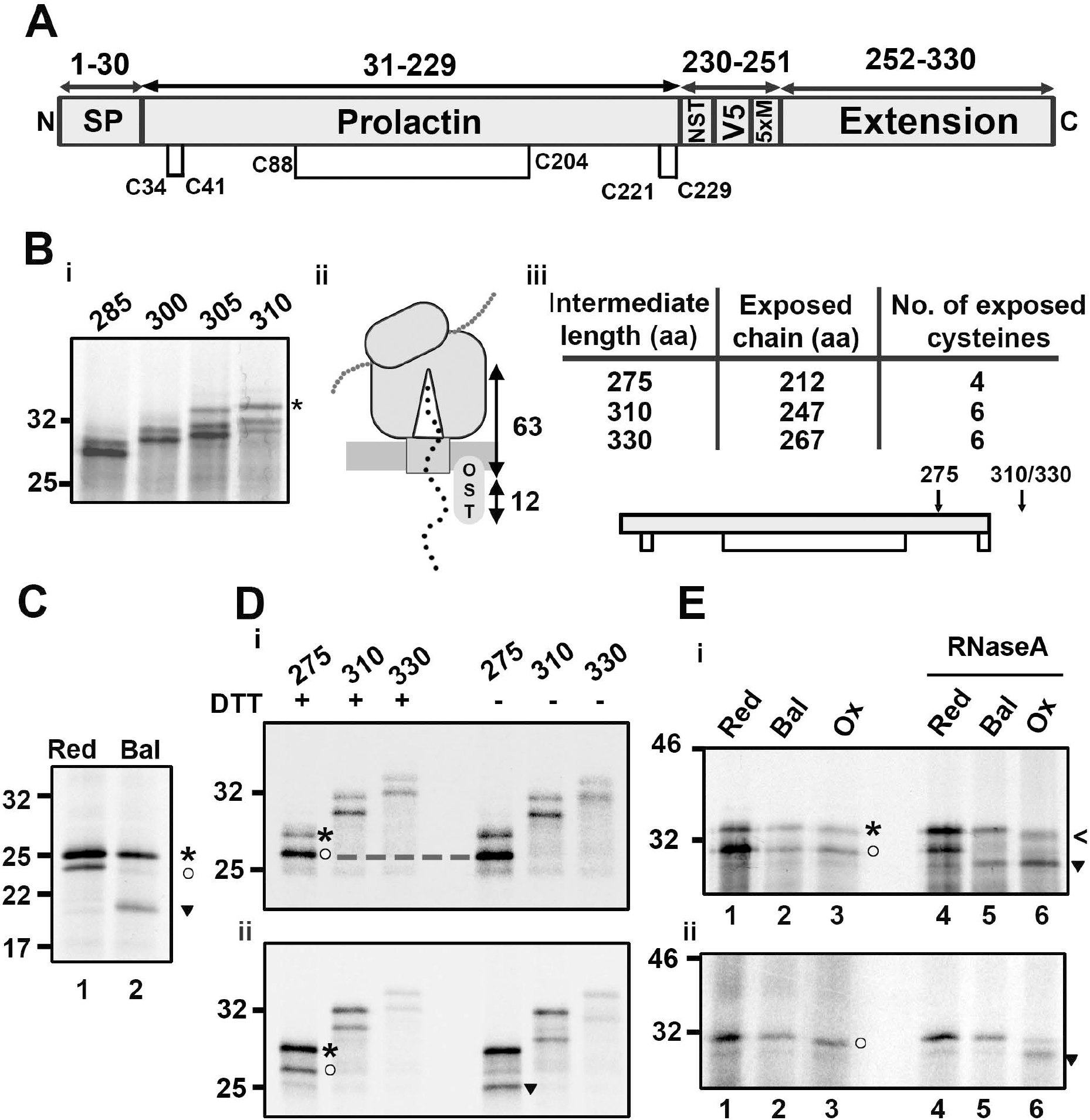
Disulfide formation in prolactin only occurs after ribosome release. (A) Organisation of the extended prolactin construct with disulfide bond positions indicated. (B) (i) Reducing SDS-PAGE of extended-prolactin intermediates AST (285-310 aa in length), translated with SP cells. Glycosylated protein is highlighted for the 310 intermediate (*), (ii) a schematic of the extension length (63 aa) required for ER-exposure and (iii) a table of prolactin intermediate lengths containing the predicted exposure of amino acids and cysteine residues to the ER-lumen and a topology diagram with ER-exposure of specific intermediates indicated by arrows. (C) Non-reducing SDS-PAGE of wild type prolactin translated with SP cells in reducing (Red) or redox-balanced lysate (Bal). (D) Extended-prolactin intermediates (275–330) translated with SP cells in redox-balanced lysate and either (i) stalled or (ii) released through RNaseA treatment. Samples were immunoisolated (V5) and run under reducing (+DTT) and non-reducing (-DTT) conditions. (E) Non-reducing SDS-PAGE of the 310 intermediate translated in reducing (Red), redox-balanced (Nat) or oxidising (Ox) lysate in (i) the presence or (ii) the absence of SP cells. Stalled samples are compared to the released (RNaseA-treated) samples. In panels C, D, E bands corresponding to reduced preprotein (*), reduced mature protein (circle), oxidised preprotein (arrow) and oxidised mature protein (triangle) are highlighted. Representative data is shown in this figure from at least 3 experimental repeats in each case.

The presence of the long-range disulfide (C88-C204) in prolactin polypeptides results in a significant mobility difference when comparing reduced and non-reduced samples on SDS-PAGE (Fig. 3C). The long-range disulfide forms in the wild type protein when synthesised in the redox-balanced lysate (lane 2). The two short-range disulfides do not influence SDS-PAGE mobility (21) and so are not detected in these assays. As prolactin contains six cysteines there is also the potential for long– range, non-native disulfides to form that may also influence gel mobility. To determine if long-range disulfides form in prolactin nascent chains we translated a range of stalled intermediates in the redox-balanced lysate and compared the resulting gel mobility under reducing (+DTT) or non-reducing (-DTT) conditions (Fig. 3Di). Of the three intermediate lengths tested (275, 310 and 330), the migration of the signal-peptide cleaved polypeptide is the same under non-reducing or reducing gel conditions, revealing an absence of disulfide formation in each case. In contrast, samples treated with RNaseA to release nascent chains from the RNCs formed disulfides (Fig. 3Dii). Based on the glycosylation results (Fig 3Biii) the 310 and 330 RNCs have a prolactin domain exposed to the ER and yet the long-range disulfide fails to form until nascent chain release from the ribosome.

We tested if the ER reductive pathway is responsible for preventing disulfide formation in the RNCs, by synthesising prolactin in the presence of SP cells in the oxidising lysate. Translation of the 310 intermediate resulted in a polypeptide with the same gel-mobility as that produced from translation in the redox-balanced or reducing lysates (Fig. 3Ei, lanes 1-3). Disulfide formation was observed only following release of the nascent chain (Fig 3Ei, lanes 5 and 6). This result demonstrates that the ER-reductive pathway is not actively reducing prolactin intermediates to prevent premature disulfide formation. We also tested whether disulfides can form in the absence of added SP-cells (Fig 3Eii). For this assay, we translated a prolactin construct that did not contain the N-terminal signal sequence (residues 31-310) and was of sufficient length (280 aa) to ensure the folding domain exited the ribosomal tunnel. When this intermediate was translated in the oxidising lysate, SDS-PAGE analysis shows an absence of disulfide formation in the RNCs (lanes 3) before RNaseA induced release (lane 6). Polypeptides synthesised in the redox-balanced and reducing lysates showed no disulfide formation in either stalled or released polypeptides. These results demonstrate that the requirement for nascent chain release before disulfide formation is independent of ER-factors.

Mechanistically, the absence of either native or non-native disulfide formation upon cysteine exposure to the oxidising milieu shows that stochastic disulfide formation is absent. This indicates that disulfide formation does not occur via the quasi-stochastic mechanism of folding, in which disulfides form first in unstructured, folding precursors before conformational folding takes place. Instead, the dependence of disulfide formation on ribosomal release is consistent with a folding-driven mechanism in which release provides the freedom that is necessary for conformational folding to occur, which then drives disulfide formation. These results thus favour the structured precursor model of oxidative folding.

### Multiple disulfide species in partially exposed disintegrin intermediates indicate a stochastic mechanism of disulfide formation

Both β2M and prolactin have stable secondary structures in which the disulfide density is low, these features may favour the structured precursor model of oxidative folding. To investigate if a contrasting protein structure folds via an alternative mechanism, we studied the disintegrin domain of ADAM10. This protein domain has a dense disulfide bonding pattern and little defined secondary structure (Fig.1 A iii) (22, 23). We illustrate the organisation of the disintegrin construct and its pattern of disulfide bonding in Fig 4A. We monitored glycosylation at position 125 to determine the intermediate length required for disintegrin domain exposure to the ER lumen (Fig. 4Bi), using the same procedure as described for the prolactin construct. Glycosylation occurred from a length of 200 amino acids onwards, which corresponds to the 63 amino acids of extension sequence required to span the length of the ribosome/Sec-channel. This is the same length observed for β2M (7) and prolactin (Fig. 3). We depict the predicted ER exposure, in terms of polypeptide length and number of cysteines exposed to the ER as a table in Fig. 4Bii, with arrows on the topology diagram illustrating ER-exposure expected at specific intermediate lengths. For the remaining experiments with disintegrin, we used a construct without the glycosylation site to simplify SDS-PAGE analysis.

**Figure 4.**
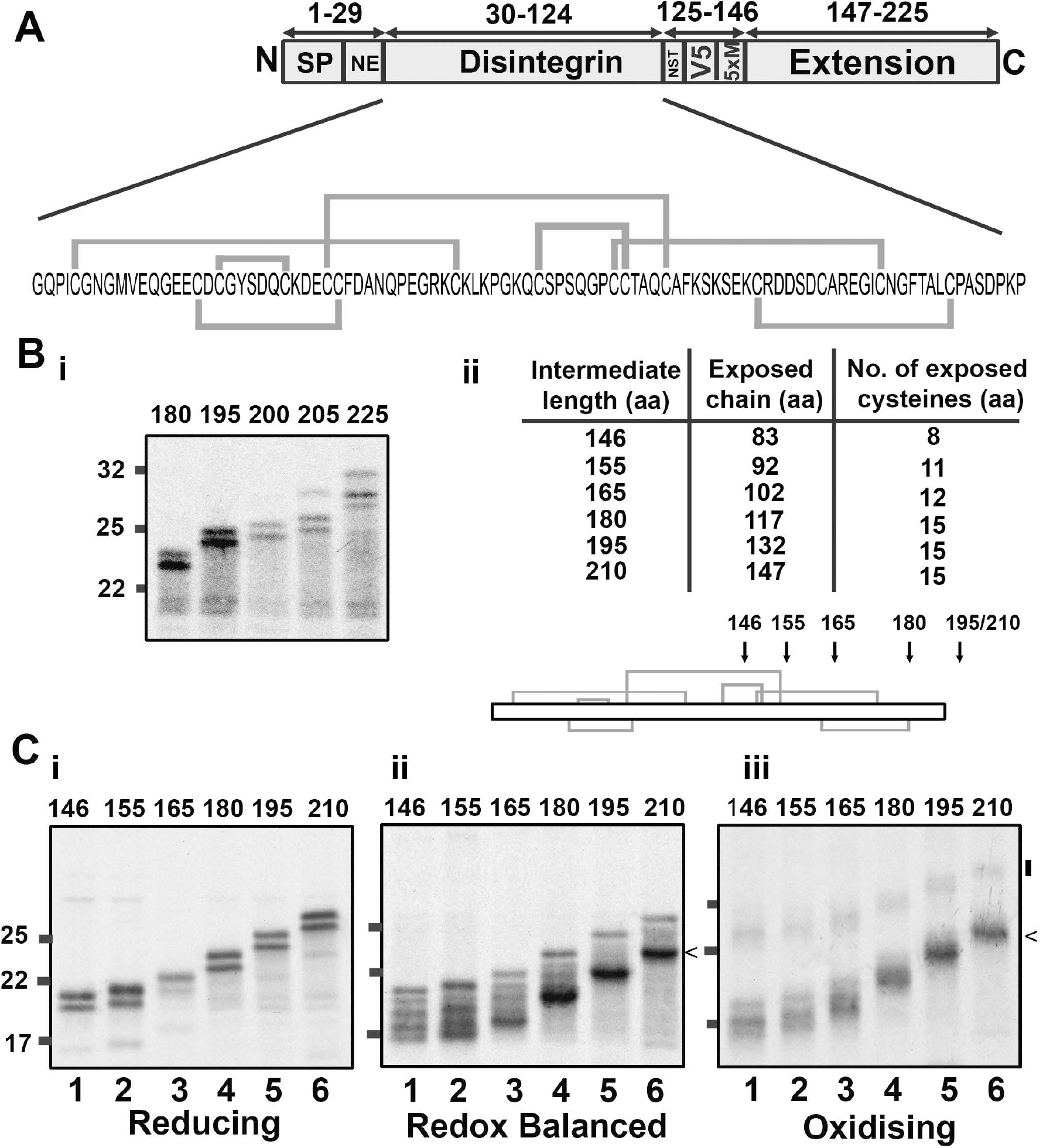
Disulfide formation occurs in a partially ER-exposed disintegrin domain. (A) Organisation of the disintegrin construct with the sequence and native disulfide bonding pattern highlighted. (B) (i) Reducing SDS-PAGE to identify glycosylation in extended disintegrin intermediates of lengths 180-225, with the table and topology diagram (ii) detailing expected exposure of amino acids and cysteine residues to the ER-lumen. (C) Non-reducing SDS-PAGE showing radiolabelled intermediates of increasing length (146-210 preprotein length) translated in (i) reducing, (ii) redox-balanced or (iii) oxidising lysate. All gels in this figure represent translation reactions with SP cells immunoisolated to the V5 epitope, and are representative of 3 independent repeats.

To evaluate disulfide bonding in the disintegrin domain relative to translocation, we expressed a range of intermediates with increasing levels of ER-exposure (Fig. 4C). When translated in the presence of SP-cells and in a reducing lysate, we observed two products in each lane that corresponded to reduced preprotein and reduced mature protein (Fig 4.Ci). When translated in a redox-balanced lysate, the shorter intermediates ran as five distinct species (Fig. 4Cii) (lanes 1-3). By comparing these to the reduced sample, we concluded that the slowest migrating species corresponds to untranslocated, reduced preprotein and that the four faster migrating species represent translocated species with different disulfide bond configurations. The clear separation of the species indicates that the disulfides produce specific long-range constraints that contribute to changes in SDS-PAGE mobility. The formation of disulfides occurs prior to full translocation of the disintegrin domain. This early disulfide formation and the mixed population of species that result indicate a quasi-stochastic mechanism of cysteine coupling. For the longer intermediates, the abundance of each gel species changes, indicating that the disulfide configuration that forms is highly dependent on the degree of ER-exposure (lanes 4-6). We observed a distinct faster migrating species once the disintegrin domain was fully exposed, (lane 6 <). This demonstrates that a single, disulfide pattern is favoured once the entire disintegrin domain enters the ER lumen.

When we preformed the same translation reactions in an oxidising lysate, a distinctly different gel-migration pattern is observed (Fig. 4Ciii). Under these conditions, disulfide formation can occur in untargeted preprotein and the ER loses the capacity for disulfide rearrangement. The absence of products that correspond to the reduced protein shows that disulfide formation is extensive. For each sample, the fast-migrating gel-species correspond to intermediates with intra-chain disulfides (< for the 210 intermediate), and the slower migrating species indicates the presence of inter-chain disulfides (vertical bar for the 210 intermediate). The intra-chain species that formed in the oxidising lysate showed more diffuse patterns on SDS-PAGE in comparison to those synthesised in the redox-balanced lysate, indicating the presence of a heterogeneous mixture of disulfide-bonded species. As the intermediates increased in length, a less heterogeneous mix of species became evident (Fig.4 C iii, lanes 5-6). These results indicate that the disintegrin disulfides formed in the oxidising lysate are slow to rearrange into distinct intermediates, resulting in an ensemble of disulfide-bonded species.

### ER-specific disulfide rearrangements occur in disintegrin nascent chains

The disulfides formed in partially exposed disintegrin intermediates are likely to be non-native disulfides that require subsequent rearrangement. Disulfide rearrangement mechanisms depend on the ER-reductive pathway. The activity of this pathway could explain why we see a different disulfide pattern when comparing partially exposed intermediates translated in redox-balanced and oxidising lysates. In the above experiments with the disintegrin domain, the translation products were immunoisolated using the V5-antibody, which recognises both preprotein and mature protein. Judging by the extent of signal-peptide cleavage in reducing samples (Fig. 4Ci), only about 50 % of the protein is ER targeted. This mixed population makes it difficult to define which of the multiple bands observed under redox-balanced and oxidising conditions originate from translocated polypeptides, and which if any originate from disulfide formation in the lysate. To define disulfide formation and rearrangements taking place in ER translocated intermediates requires a method to separate preprotein and mature protein. For this purpose we utilised a neo-epitope tag (NE) placed between the signal peptide and the disintegrin domain. This epitope is only recognised if the signal peptide is cleaved, allowing the isolation of mature protein without contamination from preprotein (24).

To identify ER specific disulfide bonded species we translated the intermediate length 146 with SP cells in reducing, redox-balanced or oxidising lysates (Fig. 5A). Samples were immunoisolated using the NE antibody or the V5 antibody and separated under non-reducing conditions. Following V5-isolation, translation products from the reducing lysate migrated as two species corresponding to preprotein and mature protein (lane 1) and a single band corresponding to mature protein following NE-isolation (lane 4). This confirms that the NE-antibody only recognises mature protein. Results with the redox-balanced lysate showed that of the five species detected following V5 isolation (lane 2) all except the reduced preprotein are ER-translocated species (lane 5). Analysis of the same intermediate translated in the oxidising lysate, showed that the fast-migrating smear is observed in both the V5-isolated (lane 3) sample and NE-isolated sample (lane 6) confirming that these products also represent ER-translocated disulfide bonded species. These results suggests that non-native disulfides form in the oxidising lysate and that the difference in disulfide bonding when comparing to the redox-balanced lysates is due to the activity of the ER reductive-pathway. The formation of non-native disulfides in the disintegrin domain indicates that cysteines are pairing through a quasi-stochastic mechanism.

**Figure 5.**
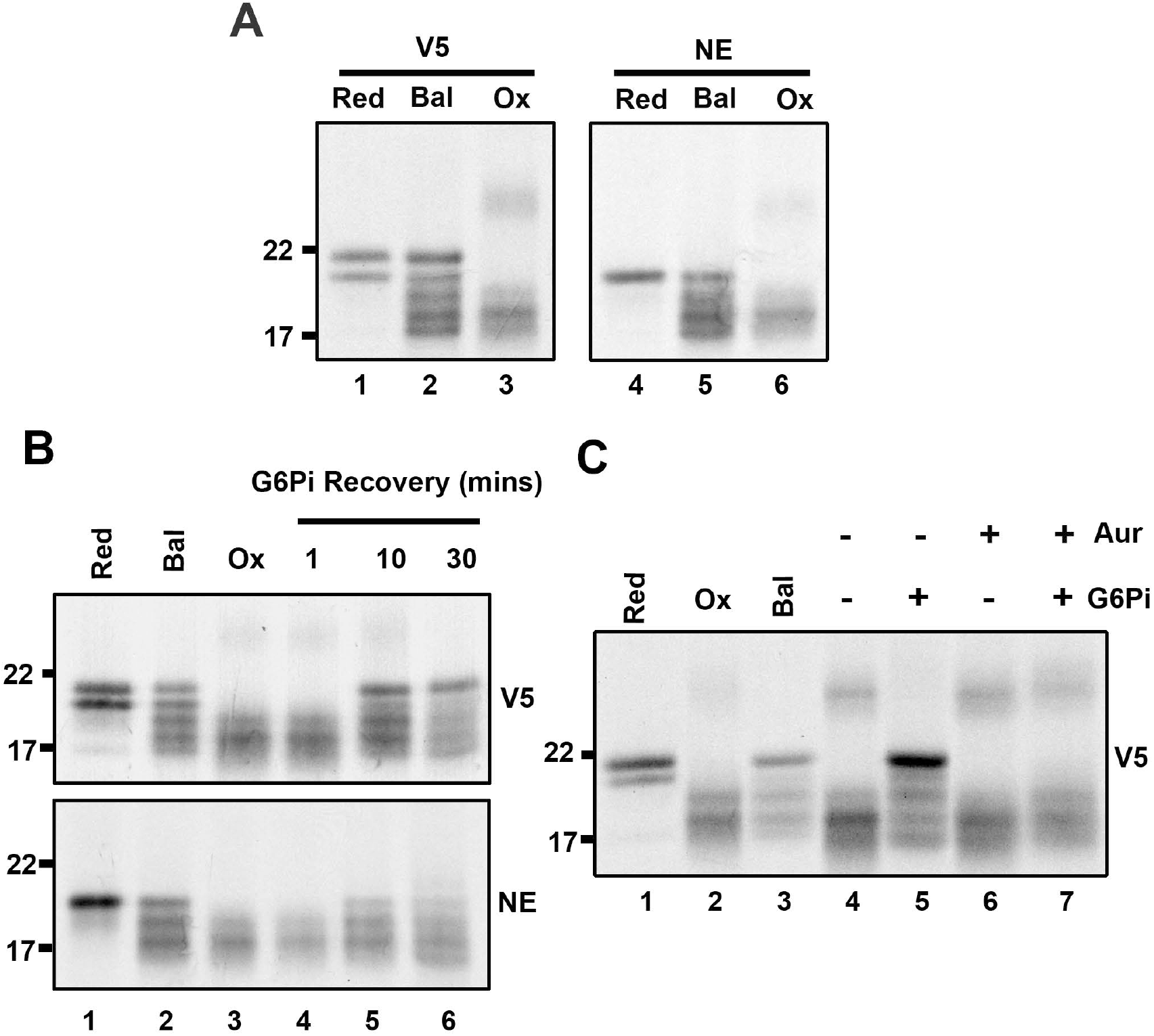
ER-specific disulfide rearrangements occur in a partially ER-exposed disintegrin intermediate. Non-reducing SDS-PAGE shows either total translation product (V5) or ER-specific species (NE) for the 146 intermediate of the disintegrin domain; (A) when translated in reducing (Red), redox-balanced (Bal) and oxidising (Ox) lysate; (B) when translated in oxidising lysate, with G6Pi added after translation and samples taken at specific time points post G6Pi treatment (lanes 4-6); and (C) when translated in oxidising lysate, to which G6Pi was added after translation in the presence or absence of auranofin (Aur) (lanes 4-7). Control samples for panels B and C were translated in reducing, redox-balanced and oxidising lysates (lanes 1-3 in both cases) for gel mobility comparison. Panel A and B were repeated 3 times and panel C twice, with representative data shown.

Having confirmed that disulfides can form in partially translocated intermediates, we next investigated if we could directly monitor disulfide rearrangements in nascent chains. To do so we synthesised the 146 intermediate in oxidising lysate and added G6Pi after nascent chain synthesis to activate the reductive pathway post-translationally (Fig. 5B). Translation products synthesised in reducing, redox-balanced or oxidising lysate were run alongside the samples for comparison (lanes 1-3). Over the 30 min time course following G6Pi addition, the mobility of the translation products changed from a diffuse smear (lane 4, 1 min) to the equivalent pattern formed in the redox-balanced lysate (lanes 5,6 10-30 min). The NE-isolated samples show the same pattern as the V5-isolated samples but without reduced preprotein. These results demonstrate that disulfide rearrangements can take place in nascent chains prior to release from the ribosome.

Finally, we confirmed that thioredoxin reductase is the source of reducing equivalents for the disulfide rearrangements observed by performing G6Pi recovery experiments in the presence of the thioredoxin reductase inhibitor auranofin. Treatment with auranofin prevented disulfide rearrangement (Fig. 5C compare lanes 5 and 7), which confirms that cytosolic thioredoxin reductase is the source of reducing equivalents for these disulfide rearrangements.

### Decreasing cysteine density of the disintegrin domain does not prevent stochastic disulfide formation

The disintegrin domain has a higher density of disulfide bonds than the substrates that follow the structured precursor mechanism. High cysteine density could be a feature that favours quasi-stochastic coupling. To test if lowering the cysteine density alters the folding mechanism we generated disintegrin constructs with specific cysteine residues substituted for serine. We designed three constructs, with 4, 3 or 1 native cysteine pairs, termed N-term cluster (4 native disulfides located at the N-terminus) triple (3 sequential disulfides) and single (1 long range disulfide). Intermediates were produced that either enable partial (146 or 165) or full (210) translocation of the disintegrin domain to the ER. Following translation, translation products was immunoisolated using V5 and Neo-epitope antibodies to compare total and ER-translocated products.

Analysis of the partially translocated N-term cluster intermediates (Fig 6Bi) showed similar gel-banding patterns to that observed for the wild type protein, with multiple species present under both balanced and oxidising conditions (lanes 2, 3 and 5, 6). This is not surprising as the ER-exposed portion of the disintegrin domain is identical to that of the wild type protein. Increasing the length of the intermediate (Fig 6Bii), resulted in a prominent fast migrating species under redox-balanced conditions (lane 8 and 11). The transition from multiple species to a single species shows that a single configuration is favoured even in the absence of the C-terminal cysteines. Synthesis of the 146 intermediate of the triple construct in the redox-balanced lysate showed two bands representing preprotein and mature protein (Fig 6. Ci, lane 2). This result contrasts to the array of disulfide-bonded species observed for the non-mutated and the N-term cluster construct and reflects the loss of specific disulfide arrangements. The preprotein runs faster in the oxidising lysate (lane 3 (o)), revealing disulfide formation in untargeted material. The ER-localised protein had a subtle change in mobility (*) when comparing the balanced and oxidising conditions (lanes 2, 3 and 5, 6) to the reduced controls (lanes 1 and 4). This indicates short-range disulfide formation occurring prior to full translocation and exposure of the domain to the ER. A more prominent disulfide-bonded species (<) is detected in redox-balanced and oxidised samples (Fig 6. C ii, lanes 8, 9 and 11, 12) upon full domain translocation, indicative of a long-range disulfide. In contrast to the non mutated protein, the process was less efficient, with the loss of cysteine residues compromising long-range disulfide formation.

**Figure 6.**
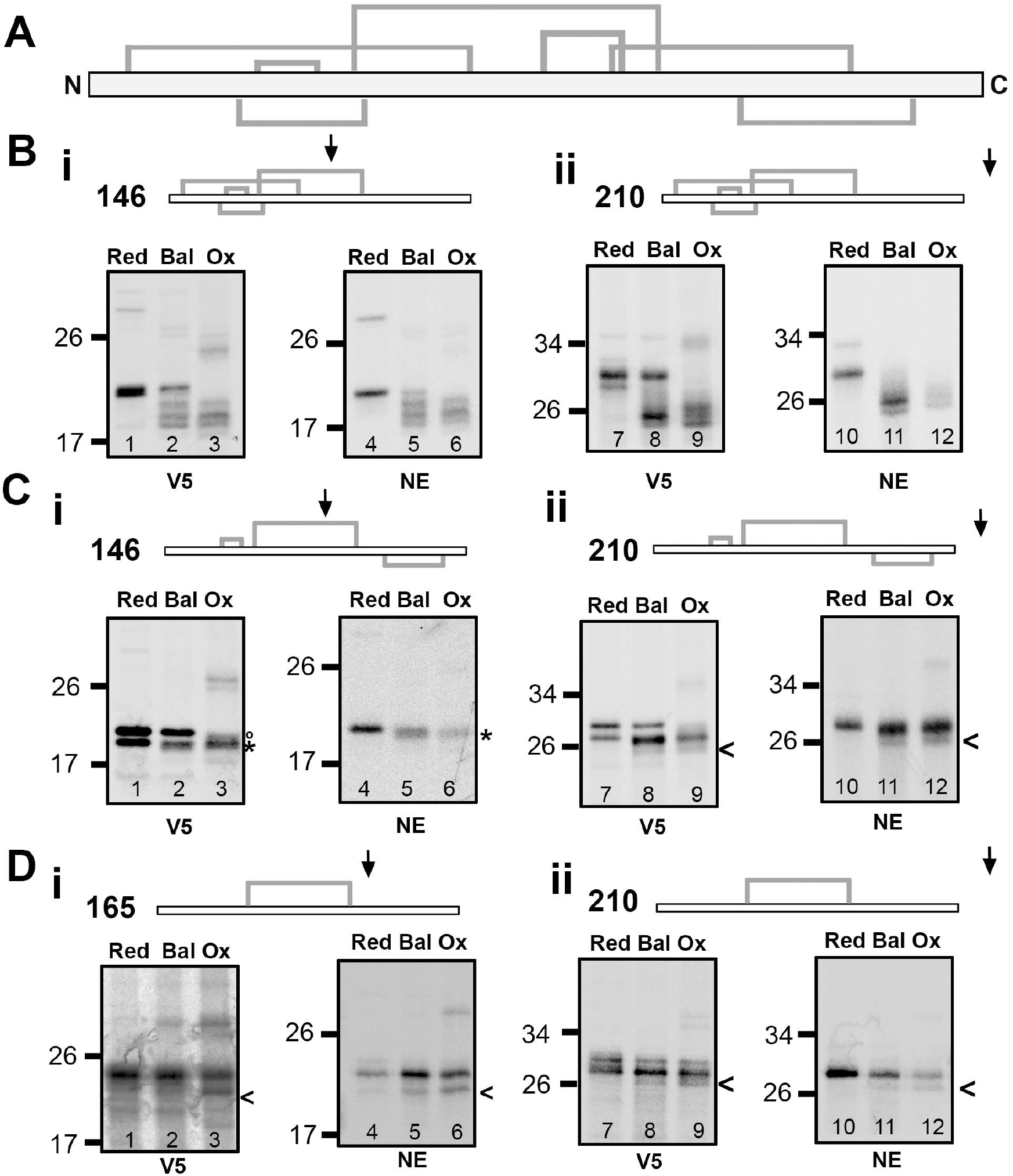
Stochastic disulfide formation occurs even when the cysteine density decreases. (A) Topology diagram showing the location of disulfides in the wild type sequence of the disintegrin domain. (B-D) Non-reducing SDS-PAGE showing disulfide formation in stalled intermediates translated under reducing (Red), redox-balanced or oxidising (ox) conditions for either (B) N-term cluster (C) triple or (D) single constructs immunoisolated using either a V5 or NE antibody as indicated. For each case (i) partially ER-exposed (146 or 165) intermediates are compared to (ii) fully exposed (210) intermediates. Topology diagrams above each panel show the position of disulfides in the relevant construct with an arrow indicating the degree of ER-exposure expected. Each condition was repeated 3 times and representative data is shown.

We translated a longer intermediate (165) for the single disulfide construct to allow ER exposure of both cysteines. Even with just two cysteines disulfide formation (Fig 6 Di (<)) occurred in the oxidising and redox-balanced lysates (lanes 2, 3 and 5, 6) prior to full domain translocation. On full translocation, the disulfide formed albeit inefficiently (lanes 8, 9 and 11, 12).

These results show that disulfide formation can occur on partial exposure of the disintegrin domain despite lowering the cysteine density. This suggests that disulfide formation remains quasi-stochastic despite the lower cysteine content. In each case, the removal of cysteines results in a less efficient disulfide formation process, indicating that disulfides in the native sequence that form stochastically in the early stages of folding influence the formation of subsequent disulfides later in the process.

## Discussion

In this study, we used an *in vitro* translation system to assess nascent chain disulfide formation, in three proteins with diverse structures and disulfide bond patterns-β2M, prolactin and the disintegrin domain of ADAM10. Our results indicate that disulfide formation occurs via two mechanisms that depend on a protein’s secondary structure. In substrates with a regular secondary structure, conformational folding drives disulfide formation. By contrast, in substrates with an atypical secondary structure, the folding of disulfide-rich domains occurs through a disulfide driven process.

For β2M, disulfide formation depends on the protein’s folding domain being fully exposed to the ER-lumen and for prolactin, the formation of the long-range disulfide requires that the protein is released from the ribosome/Sec-complex. In both cases, there is a delay in disulfide formation, despite the exposure of multiple cysteines to the ER lumen. This absence of early cysteine coupling favours the structured precursor mechanism of folding. For β2M, it is already established that disulfide formation follows folding (7) with an initial collapse of the nascent chain occurring during ER entry. In this case, the partial folding of early intermediates is likely to spatially separate cysteines and prevent disulfide formation, despite the favourable oxidising conditions. For prolactin, disulfide formation depends on the nascent chain’s release from the ribosome/Sec-complex, which suggests that the tethering of the polypeptide to the ribosome prevents folding from reaching completion. Viable explanations for this dependence on release include a requirement for C- and N-terminal interactions to initiate the folding process (25, 26), and inhibitory interactions with cellular factors (27, 28) that are alleviated on release. Although ribosome tethering clearly prevents conformational folding from being completed, the absence of cysteine coupling indicates that the translation intermediates are unlikely to be unstructured. Instead, we propose a similar mechanism to that identified for β2M in which partially folded precursors spatially separate cysteines and prevent premature disulfide formation. These findings fit with those of other studies that have demonstrated partial folding at the co-translational stage (29).

In contrast to β2M and prolactin, disulfide formation occurs in disintegrin intermediates prior to the folding domain’s full exposure to the ER lumen and results in a mixed population of non-native disulfide-bonded species. A combination of the structural restrictions on the polypeptide when it is partially ER-exposed, combined with a stochastic mechanism of cysteine coupling explains the array of species we observe. This is consistent with the quasi-stochastic model of folding where disulfides form first in an unstructured precursor before conformational folding (4). Despite this stochastic mechanism, the formation of a single disulfide bonded form on full domain exposure reveals co-operatively to the process. We conclude that the disintegrin domain has a disulfide driven mechanism of folding, where quasi-stochastic cysteine coupling in the early stages provides the structural constraints for specific cysteine coupling at the later stages.

In a previous study, we identified ADAM10 as a substrate of the reductase ERp57 along with other proteins that share densely disulfide-bonded domains with low levels of secondary structure (30). This association is evidence for non-native disulfide formation and suggests that proteins with these structural features follow a stochastic cysteine-coupling process. Further examples of proteins that form non-native disulfides during folding include LDLR (10), and Bowman-Birk inhibitor (31), both of which contain disulfide-rich, irregularly structured domains. The disintegrin constructs we investigated with lower cysteine density retained the features of the quasi-stochastic process. This suggests that secondary structure rather than the cysteine density is the deciding factor in the mechanism of cysteine coupling.

In this study, we assayed disulfide formation in stalled translation intermediates to provide insight into the folding mechanisms that achieve correct cysteine coupling in native protein structures. Our results provide novel insights into the relationship that exists between disulfide formation and protein folding. Our findings also raise questions for future studies to address, including (i) whether these mechanisms apply to a broader number and diversity of substrates; (ii) the molecular interactions involved in these folding processes; and (iii) how folding events correlate with translation in real time. Such studies will help to further characterise the mechanisms described here and will advance our understanding of the fundamentally important process of oxidative protein folding in the cell.

## Acknowledgements

This work was funded by the Wellcome Trust (grant number 103720) and the Royal Society (ref IE140492). XC is the recipient of a Chinese Scholarship Council award.

## Declaration

The authors declare that they have no conflict of interest.

## Materials and Methods

### Molecular graphics

The ribbon diagrams (Fig.1) were drawn using the UCSF Chimera package, version 1.8, from the Resource for Biocomputing, Visualization, and Informatics at the University of California, San Francisco (32) using PDB files 1A1M (β2M), 1RW5 (prolactin) and 5L0Q (ADAM10 disintegrin).

### Template generation

All protein constructs had the same C-terminal extension sequence with alternate N-terminal domains. This was either human β2M (extended-β2M), bovine prolactin (extended-prolactin) or human ADAM10 residues 456-550 (extended-disintegrin). The extended-β2M and extended-prolactin constructs contained their native N-terminal signal sequences while the extended-disintegrin construct had the human-β2M signal sequence and the Neo-epitope sequence (IIPVEEENP) added at the N-terminus. These constructs were synthesised as plasmid DNA (Table S1) using GeneArt^TM^ gene synthesis (ThermoFisher Scientific), with the sequence of each provided in supplementary material (Fig. S1–S3). Cysteine variants of the extended-disintegrin construct were also synthesised in this way (Table S1). Mutations to remove glycosylation sites were introduced by site-directed mutagenesis using ACCUZYME^TM^ (Bioline). Plasmid DNA was used as a template for PCR using an appropriate forward primer that adds a T7 promoter (Table S2) and various reverse primers that lack stop codons (Table S3). PCR products were ethanol precipitated, re-suspended in nuclease-free water, and transcribed into RNA templates using T7 RNA polymerase (Promega) (at 37 °C, 2 h). The resulting RNA was ethanol precipitated and resuspended in nuclease-free water for use as template in translation reactions. Templates containing stop codons for β2M and prolactin were generated as described previously (7, 13).

#### Cell free translation

Translations were performed using the Flexi^®^ Rabbit Reticulocyte Lysate System (Promega) supplemented with 10 mM DTT (Melford) for reducing lysate, 5 mM G6Pi (Sigma) for redox-balanced lysate or ddH20 (oxidising lysate). Typical translation reactions contained amino acids except for methionine (20 μM), KCl (40 mM), EasyTag^TM^ EXPRESS^35^S Protein Labeling Mix (PerkinElmer) (1 μl/25-μl reaction) and RNA template. Semi-permeabilised HT1080 human fibrosarcoma cells were prepared as described previously (19) and added to a concentration of ~10^5^ per 25 μl translation reaction if required. Following assembly of components, translation reactions were incubated at 30 °C for either 10 min, for glycosylation experiments and all experiments with stalled β2M and Prolactin intermediates, or 30 min, for templates with stop codons and stalled-disintegrin intermediates. After this time, samples were either treated with RNase A (Sigma) (1 mg/ml, 5 min at 30 °C) or placed directly on ice. Samples were treated with *N*-ethylmaleimide (NEM) (Sigma) at a concentration of 20 mM before further processing. Samples for glycosylation assays or downstream processing through ultracentrifugation were also treated with cycloheximide (2 mM). Alternative translation conditions were optimised for prolactin 30-310 (Fig 3. Eii), with amino acids except for cysteine replacing amino acids except for methionine and the KCl concentration increased to 50 mM.

#### Sample processing

Samples containing SP cells were centrifuged (16,000 × *g*, 30 s) following translation to isolate cell pellets, which were washed with KHM buffer and re-suspended either in SDS-PAGE sample buffer (20 μl) for direct SDS-PAGE analysis (crude samples) or in immunoisolation (IP) buffer (50 mM Tris, pH 7.5, 1% (v/v) Triton X-100, 150 mM NaCl, 2 mM EDTA, 0.5 mM PMSF, and 0.02% (w/v) sodium azide) for immunoisolation where indicated. Samples translated without SP cells, were processed by various procedures following NEM treatment. In each case, either (i) IP buffer (0.9 ml) was added for immunoisolation, (ii) crude translation product was run directly on non-reducing SDS-PAGE or (iii) ribosome-nascent chain complexes were isolated by ultracentrifugation. For the ultracentrifugation procedure, samples were loaded onto a sucrose cushion (2.5 M sucrose, 50 mM Tris pH 7.5, 10 mM MgCl_2_, 25 mM KCl) for 90 min at 265,000 x g. Pellets were rinsed with nuclease free water and re-suspended in non-reducing sample buffer for SDS-PAGE analysis.

#### Immunoisolation procedure

Samples in IP buffer were incubated with 0.5% (v/v) Protein A-Sepharose (Generon) for 30 min (4 °C) and then centrifuged (2000 × *g*, 5 min) to clear samples of material that is bound non-specifically. The resulting supernatant was incubated with Protein A-Sepharose (0.5% v/v) and the indicated antibody at 4 °C overnight. The antibodies used were the polyclonal rabbit anti-human β2M antibody (Dako) (used at 1/1000), the anti-V5 antibody (Invitrogen) (used at 1/10,000) or amino-antibody to the mature form of PLAP153 (used at 1/500) (33), which we term the Neo-epitope (NE) antibody. For NE immunoisolation procedures, PAS was pre-treated with 1% Bovine serum albumin (BSA). Following overnight incubation with the antibody, samples were isolated by centrifugation (2000 × *g*, 5 min) and washed three times with 1 ml of IP buffer and once with 100 μl of 20 mM HEPES, pH 7.5. Protein was eluted by boiling the beads in non-reducing SDS-PAGE sample buffer. If samples were to be run under reducing conditions then 10 mM DTT was added to the sample before boiling and once cooled NEM was added (1 μl, 1M NEM) before loading. Samples were run on SDS-PAGE and the gels were fixed, dried and exposed to either BioMax MR film (KODAK) for analysis by autoradiography or to phosphoimager plates for analysis on a FLA-7000 bioimager (Fujifilm).

#### G6Pi recovery and auranofin treatment

Following translation of the 146 disintegrin intermediate (10 min at 30 °C) in oxidising lysate, G6Pi was added (5 mM) and samples incubated at 30°C for 10 min. Aliquots were removed at the time points specified and treated with NEM for subsequent processing using V5 and NE immunoisolation, as described above. For the auranofin treatment, 10 minute translations were performed before auranofin (Sigma) was added at a concentration of 20 μM to relevant samples. Following a further 10 min incubation at room temperature, 5mM G6Pi was added for 30 min at 30°C. Samples were treated with NEM before V5 immunoisolation and SDS-PAGE analysis, as described above.

## Supplementary information

**Table S1:** Plasmid list

| Plasmid | Description |
| --- | --- |
| Extended-β2M | DNA Sequence in Fig.S1 |
| Extended- β2M AST | Extended- β2M with N120A mutation |
| Extended-prolactin | DNA Sequence in Fig.S2 |
| Extended-prolactin AST | Extended-prolactin with N230A mutation |
| Extended-disintegrin | DNA Sequence in Fig.S3 |
| Extended-disintegrin AST | Extended-disintegrin with N125A mutation |
| Extended-disintegrin AST: N-term cluster | Extended-disintegrin with N125A mutation and cysteines at positions 77,84,85,98,104,110 and 117 substituted for serines. |
| Extended-disintegrin AST: triple | Extended-disintegrin with N125A mutation and cysteines at positions 34, 45, 58 , 69 , 77 ,84 , 85, 104 and 110 substituted for serines. |
| Extended-disintegrin AST: single | Extended-disintegrin with N125A mutation and cysteines at positions 34, 45, 47, 53, 58, 69, 77, 84, 85, 98, 104, 110, 117 substituted for serines |

**Table S2:**
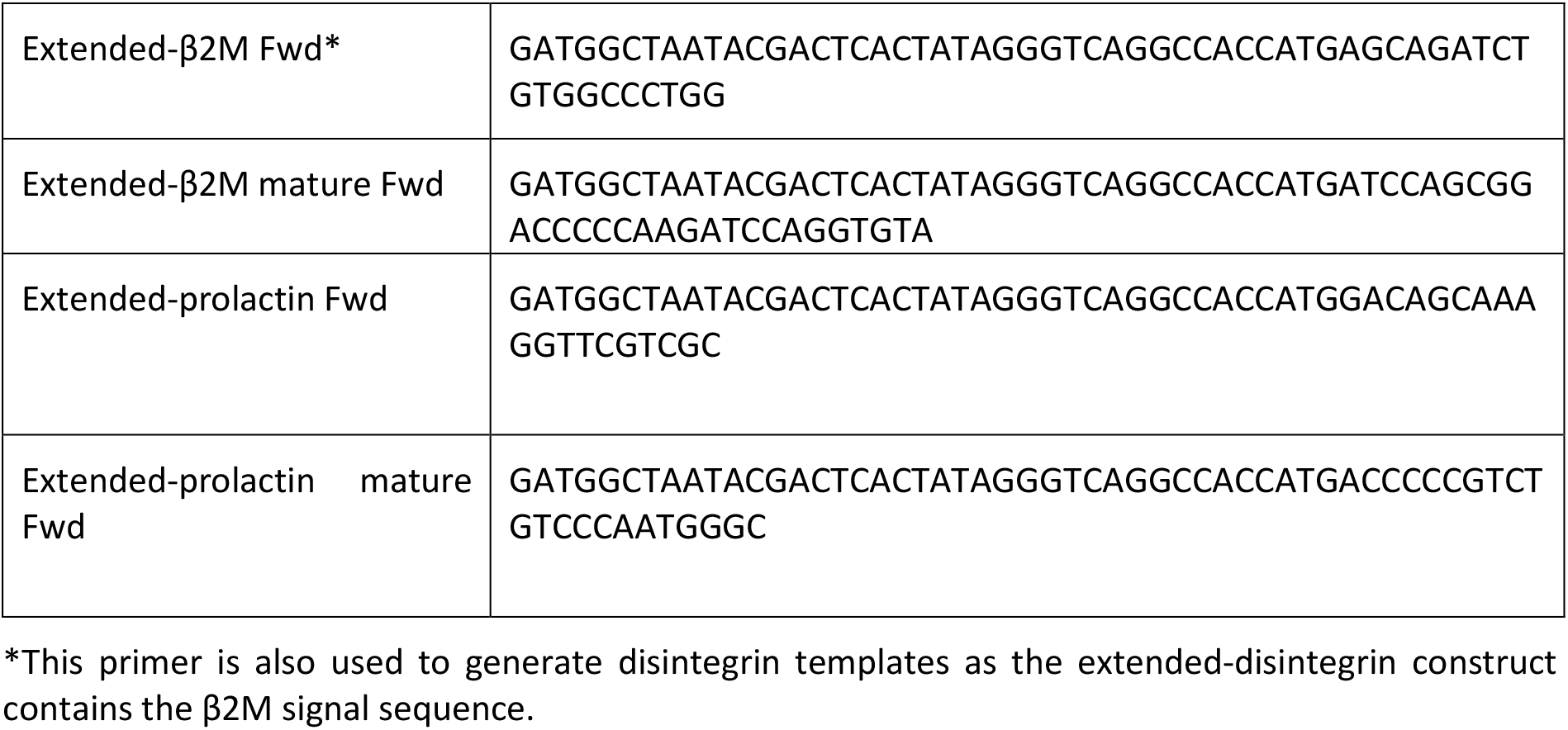
Forward primers for transcription/translation template generation

**Table S3:**
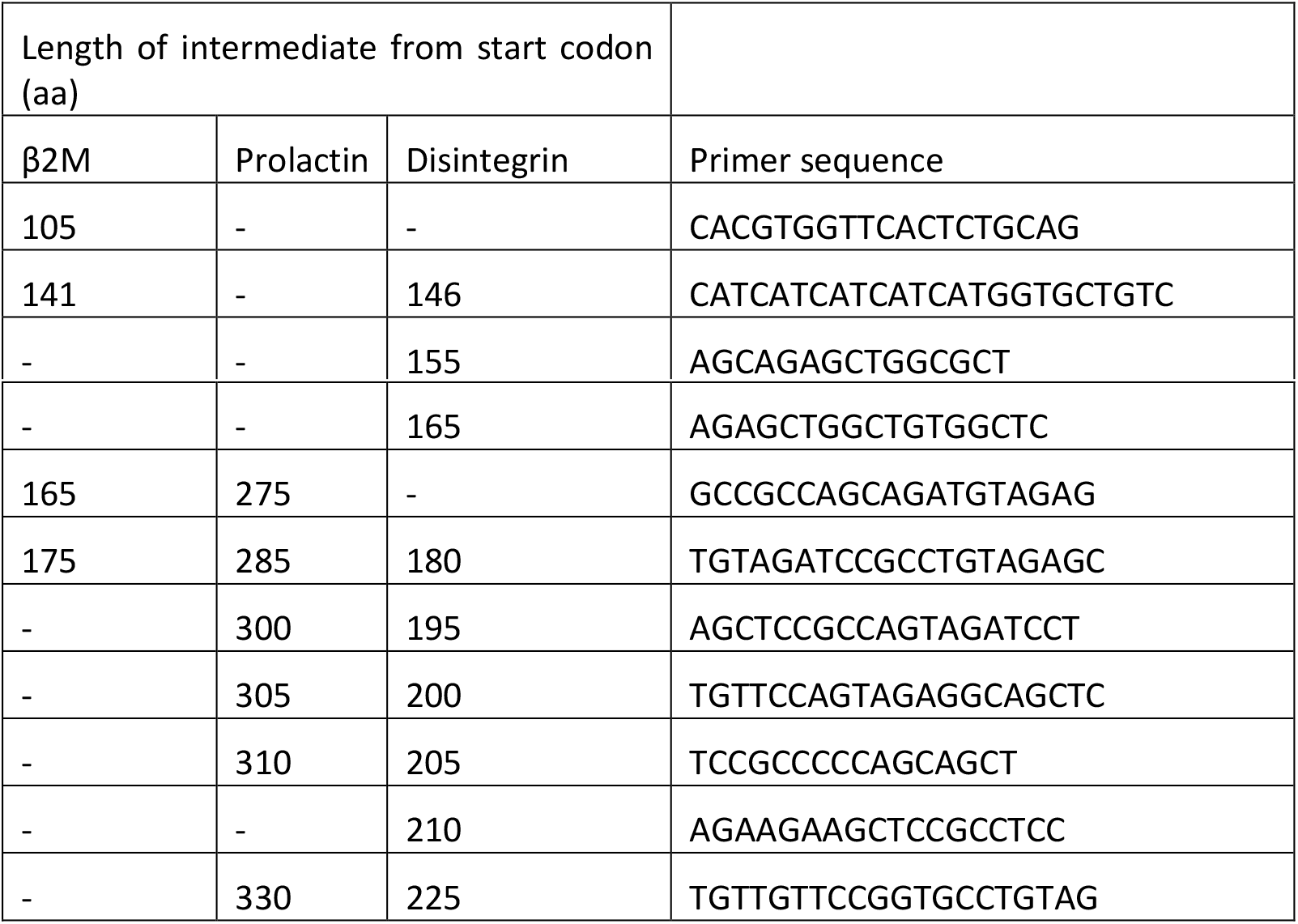
Reverse PCR primers to generate templates for stalled intermediates

**Figure S1. DNA sequence of the β2M construct**

ATGAGCAGATCTGTGGCCCTGGCTGTGCTGGCCCTGCTGTCTCTGTCTGGCCTGGAAGCCATCCAGCGGACCCCCAAGATCCAGGTGTACAGCAGACACCCCGCCGAGAACGGCAAGAGCAACTTCCTGAACTGCTACGTGTCCGGCTTCCACCCCAGCGACATCGAGGTGGACCTGCTGAAGAACGGCGAGCGGATCGAGAAGGTGGAACACAGCGACCTGAGCTTCAGCAAGGACTGGTCCTTCTACCTGCTGTACTACACCGAGTTCACCCCCACCGAGAAGGACGAGTACGCCTGCAGAGTGAACCACGTGACCCTGAGCCAGCCCAAGATCGTGAAGTGGGACCGGGACATGAACAGCACCGGCAAGCCCATCCCCAACCCTCTGCTGGGCCTGGACAGCACCATGATGATGATGATGTCCGGCACCGCCAGCGCCAGCTCTGCTGGATCTGGCGGCGGAGCCACAGCCAGCTCTACATCTGCTGGCGGCACAAGCACCGGCTCTACAGGCGGATCTACAGCAGGCGCTGCTGGCGCAACAGGCGGAGGATCTACTGGCGGAGCTGCCTCTACTGGAACAGCTGCTGGGGGCGGAGGCGGAGCTTCTTCTGGAACAGGCACAGGCGCCAGCGGCGCTACAGGCACCGGAACAACA

**Figure S2. DNA sequence of the prolactin construct**

ATGGACAGCAAAGGTTCGTCGCAGAAAGGGTCCCGCCTGCTCCTGCTGCTGGTGGTGTCAAATCTACTCTTGTGCCAGGGTGTGGTCTCCACCCCCGTCTGTCCCAATGGGCCTGGCAACTGCCAGGTATCCCTTCGAGACCTGTTTGACCGGGCAGTCATGGTGTCCCACTACATCCATGACCTCTCCTCGGAAATGTTCAACGAATTTGATAAACGGTATGCCCAGGGCAAAGGGTTCATTACCATGGCCCTCAACAGCTGCCATACCTCCTCCCTTCCTACCCCGGAAGATAAAGAACAAGCCCAACAGACCCATCATGAAGTCCTTATGAGCTTGATTCTTGGGTTGCTGCGCTCCTGGAATGACCCTCTGTATCACCTAGTCACCGAGGTACGGGGTATGAAAGGAGCCCCAGATGCTATCCTATCGAGGGCCATAGAGATTGAGGAAGAAAACAAACGACTTCTGGAAGGCATGGAGATGATATTTGGCCAGGTTATTCCTGGAGCCAAAGAGACTGAGCCCTACCCTGTGTGGTCAGGACTCCCGTCCCTGCAAACTAAGGATGAAGATGCACGTTATTCTGCTTTTTATAACCTGCTCCACTGCCTGCGCAGGGATTCAAGCAAGATTGACACTTACCTTAAGCTCCTGAATTGCAGAATCATCTACAACAACAACTGCAACAGCACCGGCAAGCCCATCCCCAACCCTCTGCTGGGCCTGGACAGCACCATGATGATGATGATGTCCGGCACCGCCAGCGCCAGCTCTGCTGGATCTGGCGGCGGAGCCACAGCCAGCTCTACATCTGCTGGCGGCACAAGCACCGGCTCTACAGGCGGATCTACAGCAGGCGCTGCTGGCGCAACAGGCGGAGGATCTACTGGCGGAGCTGCCTCTACTGGAACAGCTGCTGGGGGCGGAGGCGGAGCTTCTTCTGGAACAGGCACAGGCGCCAGCGGCGCTACAGGCACCGGAACAACA

**Figure S3. DNA sequence of the disintegrin construct**

ATGAGCAGATCTGTGGCCCTGGCTGTGCTGGCCCTGCTGTCTCTGTCTGGCCTGGAAGCCATTATTCCGGTGGAAGAAGAAAACCCGGGGCAGCCTATCTGCGGCAACGGAATGGTGGAACAGGGCGAAGAGTGCGACTGTGGCTACAGCGACCAGTGCAAGGACGAGTGCTGCTTCGATGCCAACCAGCCTGAGGGCAGAAAGTGCAAGCTGAAGCCTGGCAAGCAGTGCAGCCCTAGCCAGGGACCTTGTTGCACAGCCCAGTGTGCCTTCAAGAGCAAGAGCGAGAAGTGCCGCGACGACTCTGATTGTGCTAGAGAGGGCATCTGCAACGGCTTCACCGCTCTGTGTCCTGCCAGCGATCCTAAACCTAACAGCACCGGCAAGCCCATCCCCAACCCTCTGCTGGGCCTGGACAGCACCATGATGATGATGATGTCCGGCACCGCCAGCGCCAGCTCTGCTGGATCTGGCGGCGGAGCCACAGCCAGCTCTACATCTGCTGGCGGCACAAGCACCGGCTCTACAGGCGGATCTACAGCAGGCGCTGCTGGCGCAACAGGCGGAGGATCTACTGGCGGAGCTGCCTCTACTGGAACAGCTGCTGGGGGCGGAGGCGGAGCTTCTTCTGGAACAGGCACAGGCGCCAGCGGCGCTACAGGCACCGGAACAACA

